# Controlling for DNA dosage with Whole Genome Sequencing improves ATAC-seq peak calling

**DOI:** 10.64898/2026.04.23.720296

**Authors:** Christophe Vroland, Mohammad Salma, Artemy Zhigulev, Pelin Sahlen, Eric Soler, Laurent Bréhélin, Charles-Henri Lecellier

## Abstract

The Assay for Transposase-Accessible Chromatin using sequencing (ATAC-seq) is a scalable and sensitive method for profiling chromatin accessibility, enabling the identification of *cis*-regulatory elements (CREs) that govern gene expression in diverse cellular contexts. Although ATAC-seq is routinely applied to both bulk and single-cell samples, we reveal that its peak calling process is compromised by local biases in DNA dosage, arising not only from copy number variations (CNVs) but also from DNA replication timing (RT). These biases can distort read coverage and compromise peak detection accuracy. As part of the FANTOM consortium’s efforts to elucidate genomic regulation, we propose enhancing the MACS pipeline by integrating whole-genome sequencing (WGS) data to account for local DNA dosage effects, analogous to the use of input controls in ChIP-seq analyses. By incorporating WGS data, we demonstrate an increase in both the number and width of ATAC peaks, with improved proximity to transcription start sites (TSSs). WGS-controlled ATAC peaks exhibit canonical CRE epigenetic marks and are enriched for trait- and disease-associated genetic variants. Furthermore, the number of WGS-controlled peaks correlates more strongly with gene expression levels compared to peaks called without WGS control. Collectively, these results demonstrate that integrating WGS as a control significantly enhances the accuracy of ATAC-seq peak calling. Critically, we show that even low-depth WGS data is sufficient to improve peak calling performance, making this approach both cost-effective and readily adoptable for routine analyses. To ensure accessibility and reproducibility, we implemented this method as an open-source Nextflow pipeline. By challenging the assumption of uniform genomic visibility, our approach also holds broad implications for other DNA sequencing–based technologies.

## Introduction

The transposase-accessible chromatin using sequencing (ATAC-seq) technology has emerged as a powerful and scalable approach for genome-wide interrogation of chromatin accessibility landscapes that define cellular identity and state (17). Its protocol, which requires a limited number of input cells, makes it possible to apply ATAC-seq to scarce primary samples and heterogeneous tissues, even to single cell analysis (10). By selectively profiling regions of open chromatin without prior knowledge of the underlying epigenetic modifications or transcription factors, ATAC-seq enables the systematic annotation of cis-regulatory elements (CREs). CREs, when accessible, recruit transcription factors (TFs) (11) and host active transcription initiation (2). Delineating CREs therefore supports inference of the regulatory programs that orchestrate gene expression in both physiological and pathological contexts (23; 5; 21).

The CREs are defined as ATAC-seq peaks. While many programs exist to identify these peak regions, MACS (Model-based analysis of ChIP-seq) (50) is the most commonly used one (17). MACS was initially designed for ChIP-seq analyses and its usage for ATAC-seq requires several parameters adjustments (17). MACS uses a sliding window to identify regions that are significantly enriched relative to the genome background. For each candidate region, the expected number of reads *λ* is estimated (see below) and used to compute a p-value associated with the observed count using a Poisson distribution. Because many factors influence the local read enrichment distribution, instead of using a constant value of *λ*, MACS estimates a local value associated with the considered region. Specifically, the value of *λ*local for a specific region is defined as max(*λ*BG, *λ*1k, *λ*10k), where BG is a constant estimated from the whole genome and the *λ*x are estimated from an x kb window centered at the candidate region. ChIP-seq often uses a matched input control so the local *λ*x are estimated from that control. In contrast, ATAC-seq is frequently run without an input control and, in that case, MACS relies directly on the ATAC-seq read coverage to estimate the *λ*x. However, ATAC-seq read counts can exhibit local biases due to variations in the amount of sequenceable DNA across chromosomes, such as those caused by copy number variations (CNVs) (43; 44). Additionally, DNA replication timing (RT) further influences DNA dosage. DNA is replicated in a precise spatiotemporal program, which is local, cell-type specific and conserved in evolution, making certain DNA regions duplicated at a specific time point while others are not. As a result, depending on the number of dividing cells, early replicating DNA regions will be more abundant and available for sequencing than late replicating ones (25; 15). Hence, although the link between chromatin opening and DNA replication is fundamental to how cells faithfully duplicate their genome during cell division (32), RT may introduce DNA sequencing biases and create local heterogeneous amounts of DNA along and between chromosomes, similar to CNVs.

Here, because both CNV and RT can be inferred from whole-genome sequencing (WGS) data (7; 26), we aim to assess the impact of incorporating WGS-derived information as a control in the MACS pipeline for computing *λ*, analogously to the use of input controls in conventional ChIP-seq analyses. We observe that integrating WGS to call ATAC peaks increases both their number and their width and infers more peaks closer to Transcription Start Sites (TSSs). These WGS-controlled ATAC peaks harbour canonical CRE epigenetic marks and are enriched for genetic variants linked to particular traits and diseases. We also show that the number of WGS-controlled ATAC peaks correlates better with gene expression than ATAC peaks called in the absence of control. Our results show that incorporating WGS data into the MACS pipeline enhances the accuracy of ATAC-seq peak calling by better accounting for local DNA dosage effects. This approach, released as an open-source Nextflow pipeline, challenges the assumption that all loci have equal visibility (i.e. all genomic regions interact equally with each other) (20) and thus carries significant implications for a wide range of DNA sequencing–based methods.

## Materials and Methods

### K562 data

ATAC-seq data for K562 cells was obtained from the ENCODE Project under experiment accession ENCSR483RKN. We used the filtered BAM files from two biological replicates (ENCFF512VEZ and ENCFF987XOV), the fold change over control BigWig file (ENCFF754EAC), and pseudo-replicated peaks file (ENCFF558BLC).

For WGS data, we used paired fastq of a shotgun PE151 Illumina sequencing done on short insert fragmented gDNA from the K562 cell line was obtained from the ENCODE Project under experiment accession ENCSR711UNY (51).

Repli-seq data for S1-S6 and G2 phases were obtained from GEO dataset (GSE148362) as BigWig files (47).

CNVs were computed with Control-FREEC (8; 6) using the ENCSR025GPQ WGS data.

H3K27ac ChIP-seq data for K562 cells was obtained from the ENCODE Project under experiment accession ENCSR000AKP (51) and the pseudo-replicated peaks file ENCFF544LXB was used.

Compiled K562 STARR-seq are from (34) and are available atl https://zenodo.org/records/15079251/files/STARR-seq_catalog.tar.gz?download=1 following the path STARR-seq_catalog/starr_signal/signal_files/bigwig/k562/allENCODE_STARR_signal_K-562.bw

### MCF-7 data

ATAC-seq data for MCF-7 cells was obtained from the ENCODE Project under experiment accession ENCSR422SUG. We used the filtered BAM files from two biological replicates (ENCFF607OSL and ENCFF772EFK) and pseudo-replicated peaks file (ENCFF821OEF).

For WGS data, 101-bp paired-end reads from Illumina HiSeq from the MCF-7 cell line were obtained from SRR8652105 (paired-end fastq files: SRR8652105_1.fastq.gz and SRR8652105_2.fastq.gz) from the bioproject PRJNA523380 of the Cancer Cell Line Encyclopedia (CCLE).

### Signal correlation with CNV and replication timing

The genome was divided into non-overlapping 1000bp bins according to the Repli-seq data binning. For each bin, mean ATAC-seq signal, mean CNV value, and replication timing value were calculated using KentUtils (24) (bigWigAverageOverBed) from their respective BigWig files.

Each bin was assigned to a replication timing phase (S1-S6 and G2) based on the highest replication timing signal among the seven phases and associated to a number from 1 to 7. Spearman correlation was computed between this replication timing phase number and the mean ATAC-seq signal in each bin. Spearman correlation was also computed between the mean CNV value and the mean ATAC-seq signal in each bin.

A Support Vector Machine Regressor (SVR, from scikit-learn library with Intel Extension for Scikit-learn) was employed to predict the log-transformed mean ATAC-seq signal in each bin using standardized mean CNV and replication timing values (S1-S6 and G2 phases) as features. Hyperparameters were optimized using Ray Tune with Parzen Tree Estimator from HyperOpt on the Pearson correlation over 100 trials. Randomly selected 1% of the bins were used as a training set, and 4% of the bins were used as a validation set for hyperparameter tuning. Permutation feature importance was applied in 50 iterations using the `permutation importance` function from scikit-learn to assess the contribution of each feature to the model’s Pearson correlation coefficient.

### WGS alignment

The WGS paired fastq files were aligned to the human reference genome (hg38) using a custom Nextflow pipeline https://gite.lirmm.fr/christophe.vroland/yawgsassembly. Down-sampling was performed using seqtk https://github.com/lh3/seqtk (27). Then, by using fastp (12), quality control and adapter trimming were performed. The resulting clean fastq files were aligned to the human reference genome (hg38) using BWA-MEM (28). SAM to BAM conversion, sorting and indexing were performed using Samtools (29). Finally, duplicate marking and removal were performed using Picard MarkDuplicates https://github.com/broadinstitute/picard. BigWig files of the WGS local read depth were generated using bedtools genomecov on the resulting BAM files. Global read depth was computed by fastp on the input fastq files as the total number of bases in the input fastq files divided by the effective genome size.

### ATAC-seq peak calling with MACS3

Peaks calling was processed with a custom Nextflow pipeline https://gite.lirmm.fr/christophe.vroland/yaatacseqpeakcalling to be as comparable as possible to the ENCODE pipeline https://github.com/ENCODE-DCC/atac-seq-pipeline/tree/master described in the ATAC-seq pipeline V1 specifications document https://docs.google.com/document/d/1f0Cm4vRyDQDu0bMehHD7P7KOMxTOP-HiNoIvL1VcBt8/edit?tab=t.0. The processed BAM files for each replicate were split and merged into two pseudo-replicates by randomly assigning each read to one of the two pseudo-replicates. A merged BAM file was also created by merging the two replicates. Peaks were called on each of the three BAM files (pseudo-replicate 1, pseudo-replicate 2 and merged) using MACS3 with and without WGS as control. Peaks that were found in both pseudo-replicates and the merged BAM files were considered as the final peaks.

### Tn5 profile

Tn5 sequence preference profile file was obtained using TOBIAS (4) with default parameters on the merged bed files of ATAC-seq peaks (summits ± 500bp) called with WGS-integrating and standard pipelines as input and ENCODE DAC Exclusion List Regions as exclusion regions (ENCODE accession number ENCSR636HFF).

### ATAC-seq non-peaks

We generated a set of non-peak regions (or background regions) following the procedure described by Pampari *et al*. (37). Briefly, the entire hg38 genome was divided into overlapping bins of 2114 bp (the input length for ChromBPNet) with a 1000 bp stride. Regions that intersected with ATAC-seq peaks called by WGS-integrating pipeline and blacklisted regions (ENCODE accession number ENCSR636HFF) were excluded. From this set of candidate negative regions, we selected a number of regions (twice the size of the positive peak set) that closely matched the GC distribution of the peak regions.

### Distance to transcript starts

Transcript starts coordinates were retrieved from GENCODE V46. We defined two classes as ‘WGS-controlled’ peaks only found with WGS control (n = 58,279) and ‘standard’ peaks identified by both pipelines (n = 91,771). Because WGS-controlled peaks are wider than standards peaks, we resized all peaks considering the peak summits +/-500 bp. The two classes were defined using bedtools (40) intersect command lines:

> intersectBed -a WGS resized.bed -b std resized.bed -wa -v for ‘WGS-controlled’ peaks
>
> intersectBed -a WGS resized.bed -b std resized.bed -wa -u for ‘standard’ peaks

For each peak summit, the distance to the closest transcript start was determined using bedtools (40) closest command line

> closestBed -a resized peaks -b gencode.v46.geneStart.bed -d

The distances obtained with WGS-controlled and standard peaks were copared using R two-sided *wilcox*.*test*.

### Enrichment analyses

Fractions of resized WGS-controlled and standard peaks (see above) intersecting specific features were compared using two-sided Fisher’s exact test with Python *scipy*.*stats*.*fisher exact*.

> a=(intersectBed -a resized WGS-controlled peaks -b feature -wa -u | wc -l)
>
> b=(wc -l resized WGS-controlled peaks)
>
> c=(intersectBed -a resized standard peaks -b feature -wa -u | wc -l)
>
> d=(wc -l resized standard peaks) fisher.test([[a,c],[b-a,d-c]])

The features tested were the following:

RepeatMasker (UCSC track restricted to Alu repeats)

GWAS variants: gwas catalog v1.0-associations e109 r2023-05-07 https://www.ebi.ac.uk/gwas/docs/file-downloads

Clinvar variants downloaded 2023/06/04 https://www.ncbi.nlm.nih.gov/clinvar/ and restricted to Pathogenic and likelyPathogenic variants

GTEx eQTLS: GTEx Analysis v10 eQTL updated signif pairs https://www.gtexportal.org/home/downloads/adult-gtex/qtl

### Distribution of other peak summits around ATAC peaks

The distribution of the peak summits of H3K27ac ChIP-seq, CAGE and Remap2022 ChIP-seq data around ATAC WGS-controlled or standard peaks was determined using bedtools (40) window command line:

> windowBed -w 300 -a atac peaks.bed -b feature.narrowPeak | awk ‘if(($18-$2)/($3-$2)*<*0) print ($18-$2)/300; else if(($18-$2)/($3-$2)*>*1) print 1+($18-$3)/300; else print ($18-$2)/($3-$2)’

Precision and Recall were calculated for each feature using bedtools (40) intersect command line:

> a=(intersectBed -a atac peaks.bed -b feature.narrowPeak - wa -u | wc -l)
>
> b=(wc -l atac peaks.bed)
>
> Precision = a/b
>
> c=(intersectBed -a feature.narrowPeak -b atac peaks.bed - wa -u | wc -l)
>
> d=(wc -l feature.narrowPeak)
>
> Recall = c/d

### Correlation between the number of ATAC peaks and gene expression

Gene expression (TPM) in K562 and MCF-7 cells were downloaded from RNA-seq ENCODE data (respectively ENCFF003XKT.tsv and ENCFF721BRA.tsv). The gene annotation used in these cases was GENCODE v29. We then determined the number of ATAC WGS-controlled or standard peaks around GENCODEv29 gene starts using bedtools (40) window command line:

> windowBed -w 1000 -a gencode.v29.geneStart.bed -b WGS-controlled/standard peaks.bed | cut -f 4 | sort | uniq -c

The TPM and number of peaks obtained in K562 and MCF-7 were joined in one single file (Supplementary Table 10), which was used to compute the correlations shown in Figure 5 and Supplementary Figure 6.

### Other bioinformatic analyses

When required, hg19 coordinates were liftovered to hg38 using the UCSC liftover command line tool (19) and the hg19ToHg38.over.chain file available at this url https://hgdownload.soe.ucsc.edu/gbdb/hg19/liftOver/hg19ToHg38.over.chain.gz. PhyloP conservation scores and gnomAD genetic constraint were downloaded from the UCSC Table Browser (respectively https://hgdownload.soe.ucsc.edu/goldenPath/hg38/phyloP100way/hg38.phyloP100way.bw and hg38/gnomAD/mutConstraint/mutConstraint.bw).

### Use of artificial intelligence in preparing this manuscript

We acknowledge the use of Mistral AI’s Le Chat in preparing this manuscript, solely for refining English phrasing and providing basic editing suggestions, following the ISCB policy for acceptable use of large language models (https://www.iscb.org/about-iscb/policy-statements-bylaws-and-legal-documents/acceptable-use-of-large-language-models-policy)

## Results

### CNV and Replication Timing are confounding factors of ATAC-seq

Processed ATAC-seq signal for K562 cells was obtained from the ENCODE repository (see Materials and Methods section). K562 was selected as it is a tier-1 cell line extensively characterized across multiple epigenomic and high-throughput profiling efforts, allowing straightforward evaluation of the biological relevance of our findings. Local variations in DNA dosage were assessed by calling CNVs from WGS data. RT profiles were determined from published seven-fraction Repli-seq datasets, in which fractions S1–S6 correspond to DNA replicated during progressive stages of S-phase, and the seventh fraction (G2) represents the latest replicating regions (47). ATAC-seq signal at the example locus shown in Figure 1a is higher in copy-number–amplified regions (more than two DNA copies) and in early-replicating S1 regions, whereas it is reduced in late-replicating regions. At the genome-wide level, ATAC-seq signal is strongly correlated with copy-number state (Figure 1b) and with replication timing (Figure 1c). The *λ* parameter computed by MACS to call ATAC-seq peaks (without external control) is in fact highly correlated with both CNV and RT (Figure 1d). While CNV was previously recognized as a confounder in ATAC-seq peak calling (43; 44), our results underscore RT as an additional confounding factor. We further build a Support Vector Machine-based model, which uses CNV and RT as predictive variables and which is able to predict ATAC-seq signal with high accuracy (Spearman *ρ* correlation between predicted and observed ATAC-seq signal ∼ 0.6, Figure 1e). Features associated with RT emerged as the most important in this model (Supplementary Figure 1a). Since CNV and RT can be estimated (7; 26) and both correlate with WGS coverage (Supplementary Figure 1b and 1c), we next assessed the benefits of controlling for DNA dosage variations with WGS during ATAC-seq peak calling, analogous to input normalization in ChIP-seq analyses.

**Fig. 1.**
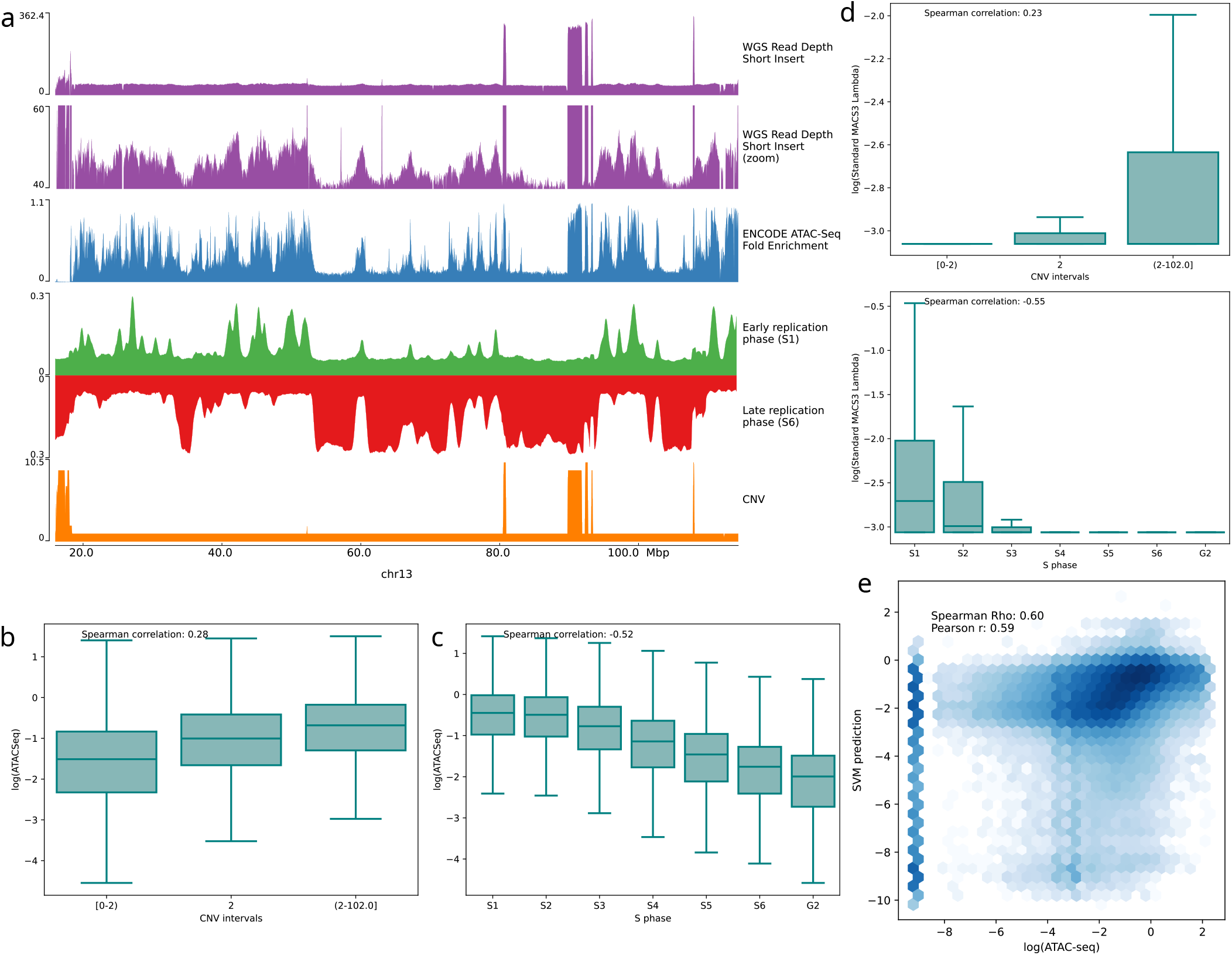
CNV and RT are confounding factors of ATAC-seq. **a** Correlation of ATAC-seq signal and whole-genome sequencing (WGS) with CNV and replication timing. A track view of ATAC-seq fold-enrichment (FE) signal (blue) and WGS local read depth (purple; top: full view, bottom: zoomed view) shows a strong positive correlation with early replication timing signal S1 (green) and CNV (orange), and an anti-correlation with late replication timing signal S6 (red), on chromosome 13. **b** Correlation of ATAC-seq signal with CNV. The Spearman correlation of ATAC-seq FE signal with mean CNV (computed in 1000-bp bins) is *ρ*= 0.28. Bins are grouped into three CNV classes (CNV *<* 2, CNV = 2, CNV *>* 2), and the log ATAC-seq FE signal is shown for each group **c**. Correlation of ATAC-seq signal with CNV and replication timing. Each bin is assigned to a replication timing phase (S1 to S6, G2) based on the highest signal among the seven replication timing tracks, and log ATAC-seq FE signal is shown per phase. The Spearman correlation with replication timing phase is *ρ* = −0.52, with higher ATAC-seq signal in early-replicating regions (S1) and lower signal in late-replicating regions (S6 and G2). **d** Correlation of MACS3 lambda signal from the standard peak-calling method with CNV and replication timing. The Spearman correlation of MACS3 lambda signal with mean CNV (1000-bp bins) is *ρ*= 0.23 (top), and with replication timing phase is *ρ*= −0.55 (bottom), with stronger signal in early phases. **e** Support Vector Machine regression model trained on CNV and replication timing (S1–S6, G2 tracks) predicts ATAC-seq FE signal. The model predicts log ATAC-seq FE signal with Spearman correlation *ρ*= 0.60.

### Controlling for DNA dosage variations in ATAC-seq peak calling

We modified the MACS peak-calling pipeline for ATAC-seq to estimate the *λ* parameter from matched WGS data, analogous to the use of input controls in standard ChIP-seq analyses (see Materials and Methods). Note that the standard peak calling pipeline used here slightly differs from the ENCODE workflow, as it employs a distinct split seed to generate pseudoreplicates and uses MACS3 instead of MACS2; yet these modifications lead to only minor differences in the resulting peaks (Supplementary Figure 2). As expected, controlling for local DNA dosage variations with WGS increased the correlation between the MACS *λ* parameter (i.e. expected number of reads) and CNV (Supplementary Figure 3a and 3b), as well as with RT (Supplementary Figure 3a and 3c). The peaks identified with WGS outnumbered peaks identified by the standard pipeline, both in terms of total peak and summit numbers, although the majority of peaks are found by both pipelines (Figure 2a and Supplementary Table 1). Narrowpeaks called with WGS (n = 150,050) were nearly twice as wide as Narrowpeaks called with a standard pipeline (n = 117,397, Figure 2b and Supplementary Table 2). To further compare peaks identified with and without WGS, we defined two classes: ‘WGS-controlled’ peaks only found with WGS control (n = 58,279) and ‘standard’ peaks identified by both pipelines (n = 91,771, Figure 2a and Supplementary Table 1). While standard peaks were enriched at annotated TSSs, WGS-controlled peaks were predominantly localized in their immediate vicinity (Figure 2c). The ATAC-seq signal in WGS-controlled peaks was slightly lower than that observed in standard peaks (Figure 2d and Supplementary Table 4), even when correcting for Tn5 sequence bias computed by the ATACorrect approach (3) (Supplementary Figure 4). Together, our findings indicate that the standard MACS pipeline for ATAC-seq peak calling, lacking a matched control, masks many open chromatin regions. This raises the question of whether the novel peaks identified through WGS integration correspond to biologically relevant regions.

**Fig. 2.**
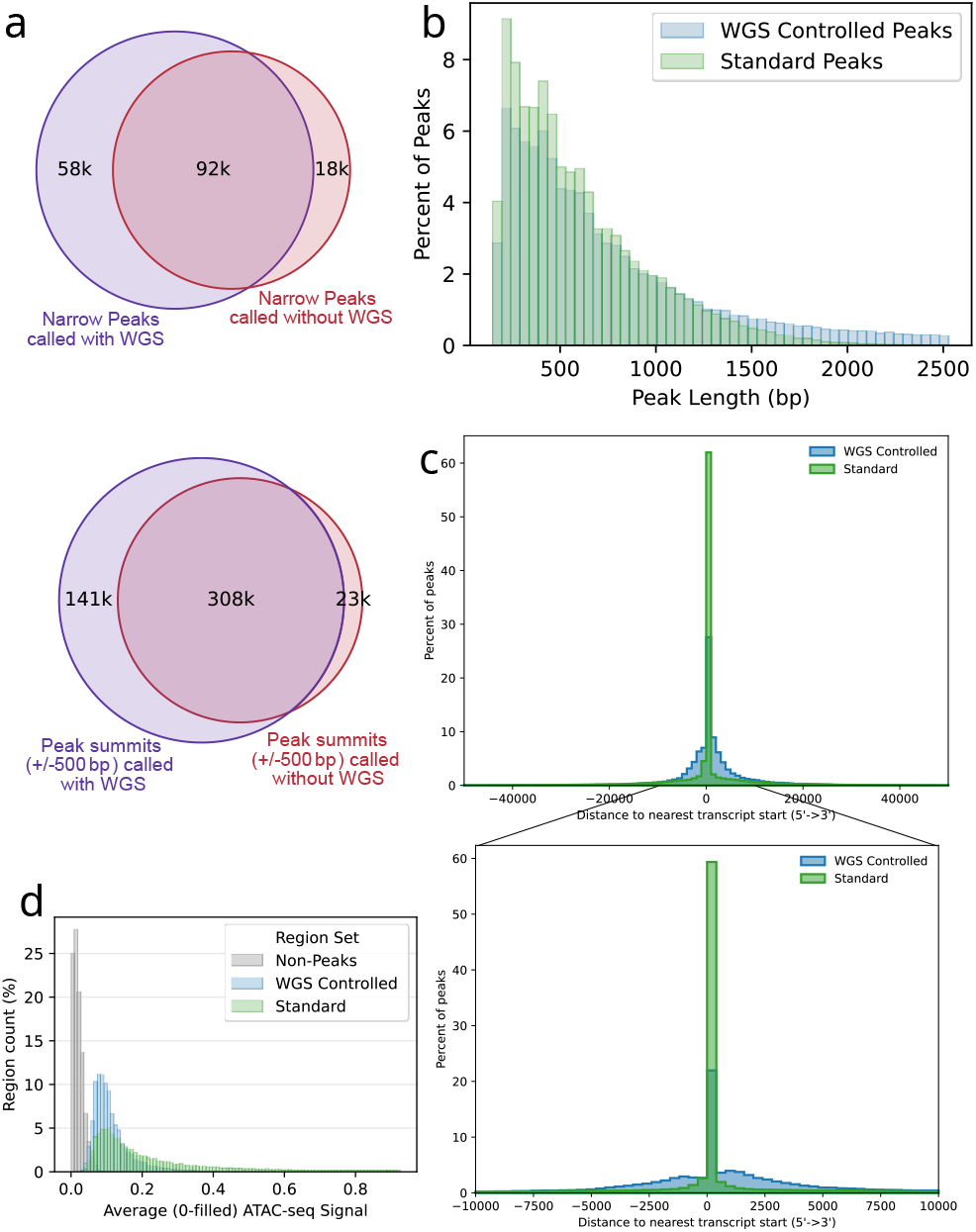
WGS-controlled peaks harbour specific features. **a** Peak calling with WGS control identifies more peaks and peak summits than the standard method. Integrating WGS in peak calling identifies 448,359 peak summits in 150,050 peak regions, compared with 230,870 peak summits in 117,397 peak regions with the standard method. A total of 91,771 peaks called with WGS intersect 99,142 peaks called with the standard pipeline, and 307,699 peak summits called with WGS (±500 bp) intersect 208,200 peak summits called with the standard pipeline. **b** Peak lengths of WGS-controlled vs. standard peaks. mean/median length = 624/496 bp for WGS-controlled peaks; mean/median length = 376/334 bp for standard peaks. **c** Distribution of peak summit distance to nearest transcript start (position 0). **d** Histograms of ATAC-seq FE signal in WGS-controlled and standard peak summits. Mean ATAC-seq FE signal was computed in a ±500 bp window around peak summits. WGS-control-only summits show lower FE signal (mean 0.118) than intersecting summits (mean 0.382). ATAC-seq signal was also compared to that observed in non-peak regions (see Materials and Methods section).

### Using WGS as control improves ATAC-seq peak calling

We first observed that WGS-controlled peaks are significantly enriched for Alu elements (Supplementary Figure 5a and Supplementary Table 3), which contribute to enhancer–promoter specificity (31). WGS-controlled peaks contain expression quantitative trait loci (eQTLs) even at a slightly higher proportion than standard peaks (Supplementary Figure 5b and Supplementary Table 3). We then examined orthogonal datasets generated in K562 cells. First, we looked at H3K27ac, which serves as a hallmark of active enhancers (41). At the representative locus shown in Figure 3a, the H3K27ac signal aligned more closely with WGS-controlled peaks than with standard peaks. At the genome scale, H3K27ac peak summits were more frequently encompassed within WGS-controlled peaks than within standard peaks (Figure 3b). Similar observations were made for peak summits from ReMap TF ChIP-seq (18) and CAGE (1) datasets (Figure 3b). Additionally, we analyzed a compilation of STARR-seq data generated in K562 cells (34). Our findings indicate that, following the ATAC-seq signal, the STARR-seq signal, which measures enhancer activity through plasmid-based reporter assays, was slightly reduced in WGS-controlled peaks relative to standard peaks, yet remained elevated compared to the signal observed in non-peak regions (Figure 3c and Supplementary Table 5). The WGS-controlled peaks were also found enriched in GWAS phenotypical traits (Figure 4a and Supplementary Table 6) and in Clinvar disease-associated pathogenic variants (Figure 4b and Supplementary Table 6). Using mutational constraint metrics from gnomAD (13) and phyloP conservation scores (38), we showed that inter-individual variations and inter-species conservation were comparable between WGS-controlled and standard peaks (Figure 4c and Supplementary Tables 7 and 8).

**Fig. 3.**
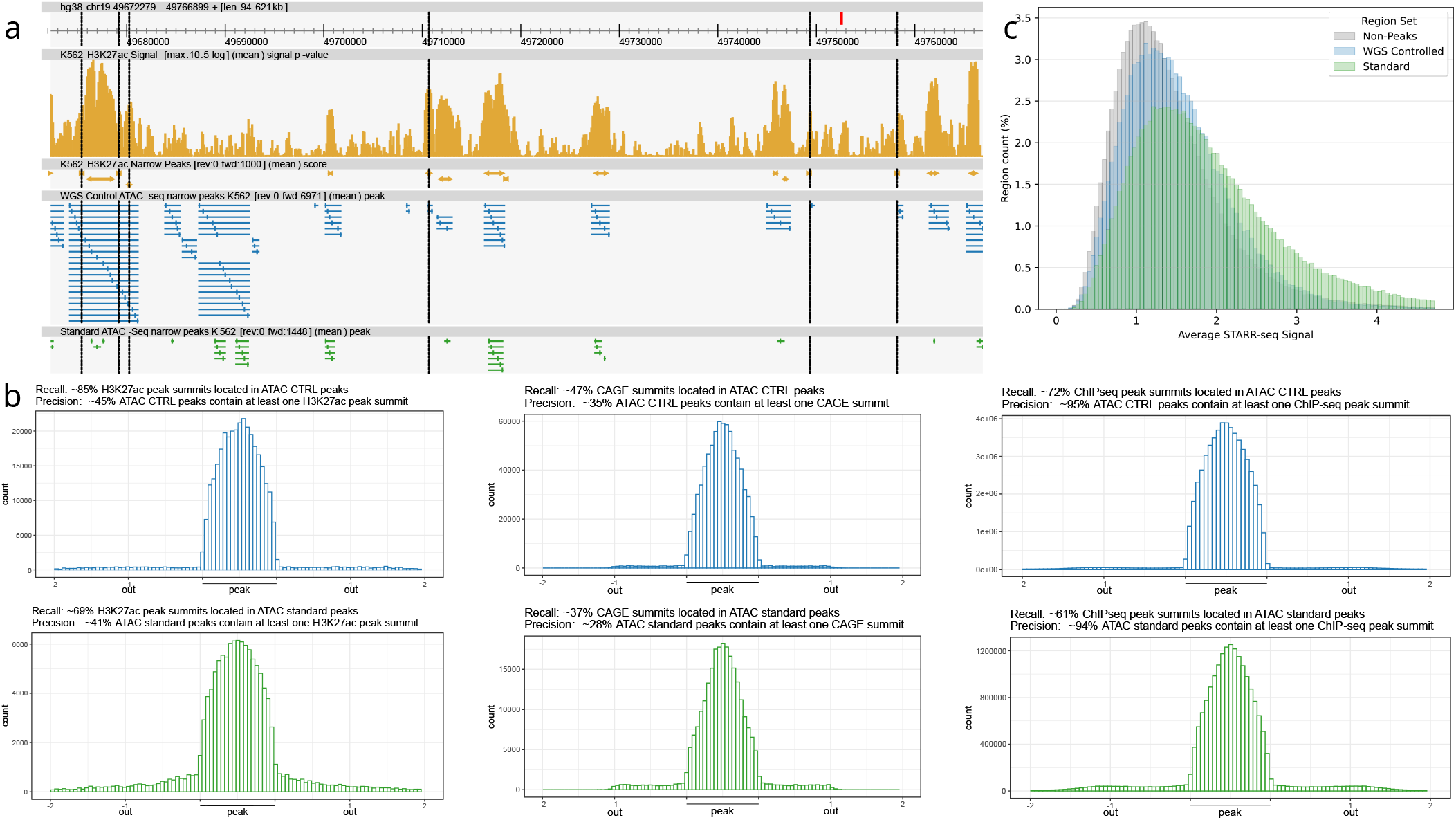
Using WGS as control improves ATAC-seq peak calling. **a** Track view of ATAC-seq peaks called with WGS-control and standard methods (chr19:49,672,279–49,766,899). From top to bottom, first track: H3K27ac peaks from ENCODE ENCSR000AKP (ENCFF544LXB); second track: ATAC-seq peaks called with WGS control; third track: ATAC-seq peaks called with standard pipeline. In ATAC-seq tracks, each peak summit is shown within its peak region as vertical line. Vertical black dashed lines indicate positions where an H3K27ac peak intersects a WGS-controlled peaks but not with standard ones. **b** Distribution of H3K27ac peak summits (left), CAGE peaks (center), and transcription factor binding sites (TFBS; right) in and around WGS-controlled (top, blue) and standard peaks (bottom, green). To compare peaks with different widths, each peak is scaled between 0 and 1 (relative start to end). Flanking regions are fixed 300bp-long windows. Recall is the proportion of H3K27ac summits, CAGE peaks, or TFBS located within ATAC-seq peaks. Precision is the proportion of ATAC-seq peaks containing at least one H3K27ac summit, CAGE peak, or TFBS. WGS-controlled peaks show higher recall and precision than standard peaks for H3K27ac (recall 85% vs 69%; precision 45% vs 41%), CAGE (recall 47% vs 37%; precision 35% vs 28%), and TFBS (recall 72% vs 61%; precision 95% vs 94%). **c** Histograms of STARR-seq signal in WGS-controlled and standard peak summits. Signal is computed in a ±500 bp window around summits with minimum coverage 0.7. STARR-seq signal was also compared to that observed in non-peak regions (see Materials and Methods section).

**Fig. 4.**
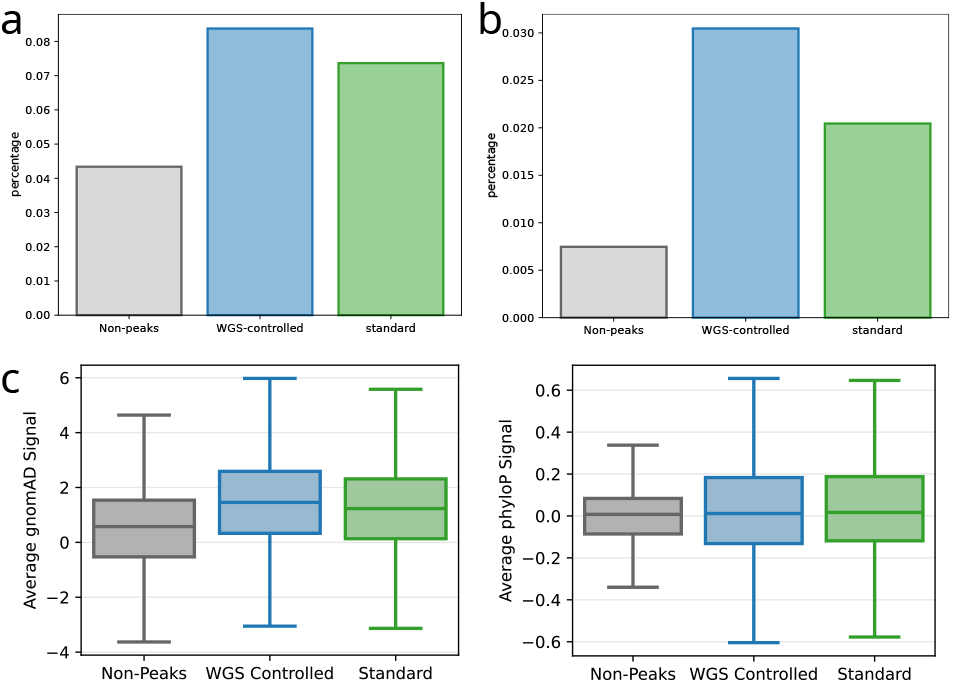
Genetic variations and conservation in WGS-controlled peaks. **a** Percentage of peaks (summit ±500 bp) overlapping GWAS causal variants. WGS-controlled peaks are more enriched than standard peaks (8.4% vs 7.4%; Fisher exact test *p* = 1.34 × 10^−31^). Percentage of non-peak regions (grey, see Materials and Methods section) is also shown for comparison. **b** Percentage of peaks (summit ±500 bp) overlapping ClinVar pathogenic variants. WGS-controlled peaks are more enriched than standard peaks (85.3% vs 82.4%; Fisher exact test *p* = 6.60×10^−133^). Percentage of non-peak regions (grey, see Materials and Methods section) is also shown for comparison. **c** GnomAD (left) and phyloP (right) scores in WGS-controlled (blue) versus standard (green) peaks. Scores are computed in a ±500 bp window around summits with minimum coverage 0.7. Both groups show similar score distributions. Scores in non-peak regions (grey, see Materials and Methods section) is also shown for comparison.

**Fig. 5.**
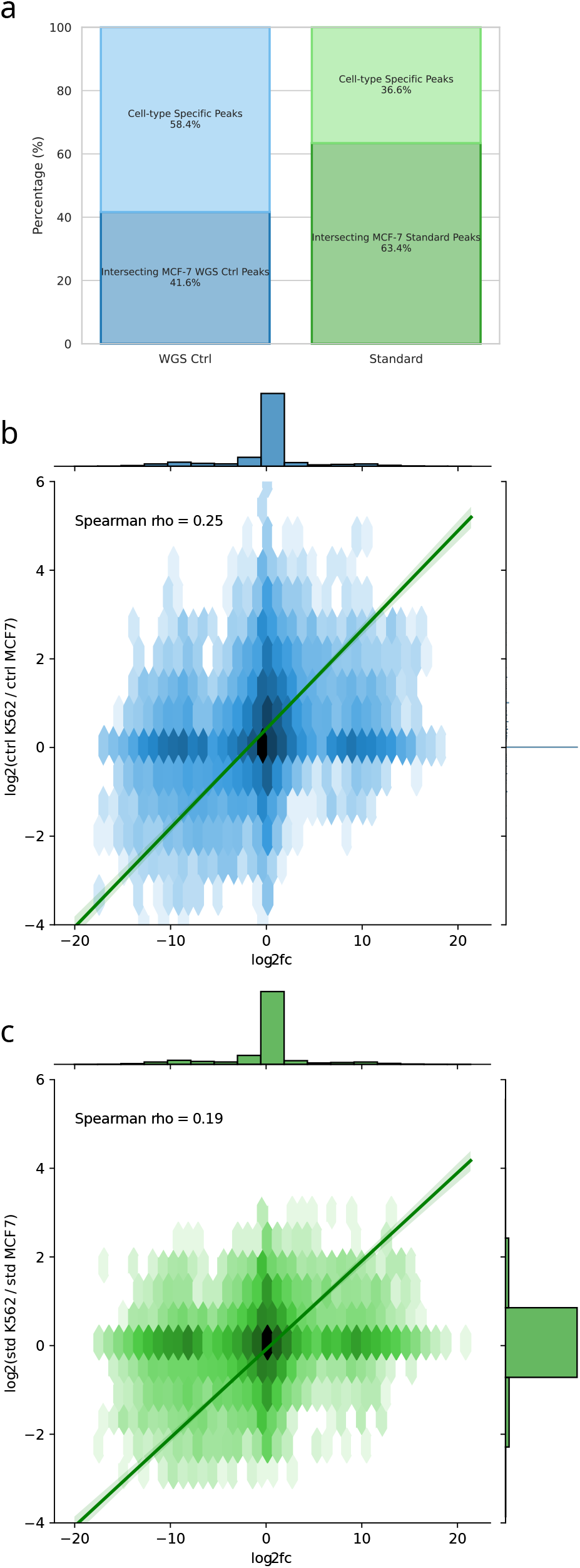
WGS-controlled peaks are informative for gene expression. **a** Fraction of WGS-controlled peaks called in K562 intersecting with MCF-7 WGS-controlled peaks and fraction of standard peaks called in K562 intersecting with MCF-7 standard peaks (respectively 41.6% vs 63.4%; Fisher exact test *p* = 0). Peaks correspond to summit ±500 bp. **b** Hexbin plots showing the log fold change of the number of peaks called with the WGS-integrating pipeline within ±500 bp of gene starts (y-axis) and log fold change of gene expression in K562 vs. MCF-7 cells (x-axis). Spearman *ρ* = 0.25. c Same as b, using peaks called with hte standard pipeline in K562 and MCF-7 cells. Spearman *ρ* = 0.19

### Integrating WGS as control reveals cell-specific ATAC-seq peaks

When comparing peak calling between K562 and MCF-7 cells, we observed that WGS-controlled peaks are more cell-specific than standard peaks: only 41.6% of K562 WGS-controlled peaks were detected in MCF-7 cells, compared to 63.4% of standard peaks (Figure 5a and Supplementary Table 9). Because the number of open chromatin regions around a gene is strongly associated with its expression level (5), we computed the correlation between the gene expression (as measured by RNA-seq) and the number of ATAC-seq peaks called with the WGS-integrating pipeline and located within a 2-kb window centered on gene start sites. This correlation was similar to, yet marginally higher than, that obtained using peaks from the standard pipeline (compare Supplementary Figures 6a and 6b). We then tested whether peaks called with WGS better predicted differential gene expression. We calculated the ratio of WGS-called ATAC-seq peaks around gene start sites between K562 and MCF-7 cells and correlated this ratio with the fold change in RNA-seq counts (K562/MCF-7). In that case, correlation using peaks called with WGS substantially exceeded that using peaks called with standard pipeline (compare Figures 5b and 5c), in line with the distribution of WGS-controlled peaks around TSSs (Figure 2c).

### Low-depth WGS is suffcient to enhance ATAC-seq peak calling

Finally, to determine the required sequencing depth for this improvement, we down-sampled the WGS data (see Materials and Methods) and observed that a depth of 5 suffices to recover the peaks used in the previous analyses (Supplementary Figure 7). This sequencing depth demonstrates that incorporating a WGS control sample into ATAC-seq analysis is cost-effective and warrants routine implementation.

## Discussion

Our findings demonstrate that ATAC-seq signals exhibit a robust correlation not only with CNVs (43; 44) but also with RT. While previous studies have documented the association between ATAC-seq and RT, with early-replicating regions exhibiting higher ATAC-seq accessibility and late-replicating regions showing reduced accessibility (45), the potential confounding effect of this relationship had not been envisaged. To elucidate the impact of DNA dosage variations on ATAC-seq data analysis, we examined their influence on background estimation within the MACS peak calling pipeline. To address this, we integrated information from WGS as a control, drawing an analogy to the use of input samples in conventional ChIP-seq analyses. Incorporating whole-genome sequencing (WGS) as a control revealed substantial deviations from results obtained using the standard MACS pipeline, which lacks matched input data. Peaks identified with WGS were not only more numerous but also broader and more frequently localized near TSSs. These WGS-controlled peaks significantly improved the prediction of differential gene expression, with potential implications for enhancer-based gene regulatory networks (eGRNs). Thus, accounting for DNA dosage can unmask regulatory regions that might otherwise remain undetected. Complementary analyses using orthogonal datasets further supported the biological relevance of the newly identified peaks and their distinct properties. Collectively, our results advocate for an improved ATAC-seq peak calling strategy that explicitly integrates DNA dosage information. While our work demonstrates the utility of incorporating WGS as input control, we do not exclude that alternative or composite control models could prove equally or even more effective. For instance, integrating GC content and RT profiles directly from Repli-seq with WGS data may refine background normalization further. Indeed our results indicate that RT significantly affects peak calling, particularly in highly proliferative cell lines such as K562 and MCF-7. Given that RT is a temporal and cell–specific feature, these findings underscore the importance of considering replication dynamics when comparing chromatin accessibility across heterogeneous or cancer-derived cell types (14). The relationship between DNA dosage and chromatin accessibility also underlies the feasibility of inferring CNVs from ATAC-seq data (44).

Although bulk and single-cell ATAC-seq differ substantially in input requirements, data sparsity, and analytical scope, most single-cell workflows continue to rely on aggregated read profiles followed by MACS-based consensus peak calling (36). This methodological continuity implies that the confounding effects described in the present study are likely to extend to single-cell ATAC-seq analyses. Furthermore, analogous to the inference of CNV from scATAC-seq data (44), RT can also be recovered from such data (22). Accordingly, provided that low-coverage WGS data are sufficient (Supplementary Figure 5), future methodological developments may benefit from incorporating matched single-cell WGS datasets (36), thereby offering a direct counterpart to the bulk WGS controls proposed herein.

Our results on ATAC-seq suggest that RT and CNV may act as pervasive confounders across a broad spectrum of genomic technologies based on DNA sequencing. In particular, several technologies able to detect all-to-all RNA-DNA interactions have been developed with varying technical details (30; 48; 9; 16) and different computational analysis methods (30; 48; 9; 16; 35; 49). Although these methods differ in the statistical tests used, they all correct for confounding factors centered on RNA, such as spurious trans-chromosomal mRNA-DNA interactions (30; 35), RNA abundance (49) and distance between the RNA-emitting gene and RNA-receiving DNA region (35; 49). We observed however that CNV and RT are also highly correlated to RADICL-seq interaction counts (46), suggesting that these features may persist as significant confounders even after accounting for RNA-specific corrections. Likewise, Hi-C normalization methods, such as iterative correction, typically rely on the assumption of equal locus visibility, which ensures that each genomic locus contributes equally to the total interaction signal and facilitates comparability across the genome (20). However, this assumption may not always be valid, particularly in the context of CNVs. In such cases, amplified genomic regions are more likely to be captured during library preparation, while regions with lower copy numbers may be underrepresented (42). This limitation likely extends to scenarios involving DNA replication, where RT can introduce additional biases as shown in this study with ATAC-seq data. Given the strong correlations observed between Hi-C contact frequencies and replication timing, it may be necessary to refine Hi-C normalization approaches to better account for replication-related biases. Addressing these challenges will not only improve data accuracy but also deepen our understanding of the mechanistic links between DNA replication, chromatin organization and the formation of topologically associating domains (33; 39).

## Data availability

The data underlying this article are available in [repository name, eg, the GenBank Nucleotide Database] at [URL], and can be accessed with [unique identifier, eg, accession number, deposition number].

## Author contributions statement

C.V., L.B. and C-H.L. conceived the scientific strategy. C.V., M.S., E.S., L.B. and C-H.L. analyzed and interpreted data. A.Z and P.S. computed CNVs in K562. C.V., M.S. and C-H.L. conducted the experiments. E.S., L.B. and C-H.L. acquired funding. C.V. and C-H.L. wrote the manuscript. All authors have read and approved the manuscript.

## Acknowledgments

We thank Anthony Mathelier, Vipin Kumar, Jean-Christophe Mouren, Benoît Ballester, Pierre-François Roux, Laurent Le Cam and all members of the IGMM/LIRMM/IMAG Computational Regulatory Genomics team for insightful discussions, data sharing and/or critical readings of the manuscript. We are indebted to the researchers around the globe who generated experimental data used in this work and made them freely available.

## Supplementary figures

**Fig. S1:**
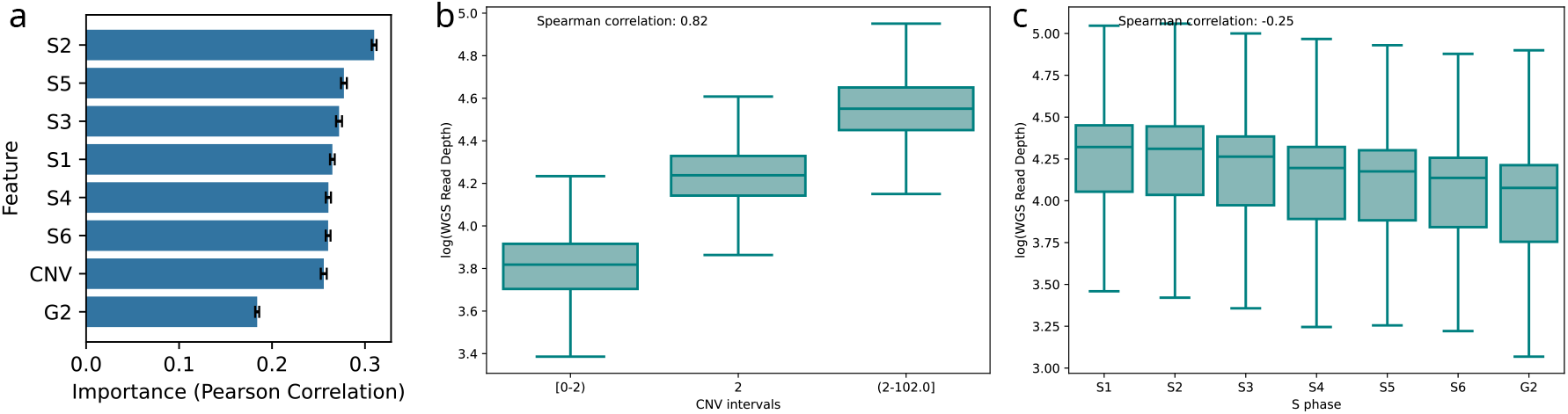
**a** Feature importance of the SVM model trained on CNV and replication timing (S1–S6, G2 tracks) to predicts ATAC-seq FE signal. **b** Boxplots of log-transformed WGS signal (y-axis) versus CNV (x-axis). Spearman correlation was computed between mean CNV and mean log WGS read depth in 100-kb bins. For boxplots, CNV is grouped into CNV *<* 2, CNV = 2, and CNV *>* 2. WGS signal is positively correlated with CNV (*ρ* = 0.82). WGS local read depth was computed using `bedtools genomecov` on the WGS BAM file. **c** Boxplots of log-transformed WGS signal (y-axis) versus replication timing (x-axis). The replication timing phase (S1 to S6, G2) with the maximum signal is assigned to each bin. WGS signal is negatively correlated with replication timing phase, with higher signal in early phases and lower signal in late phases (*ρ* = − 0.25).

**Fig. S2:**
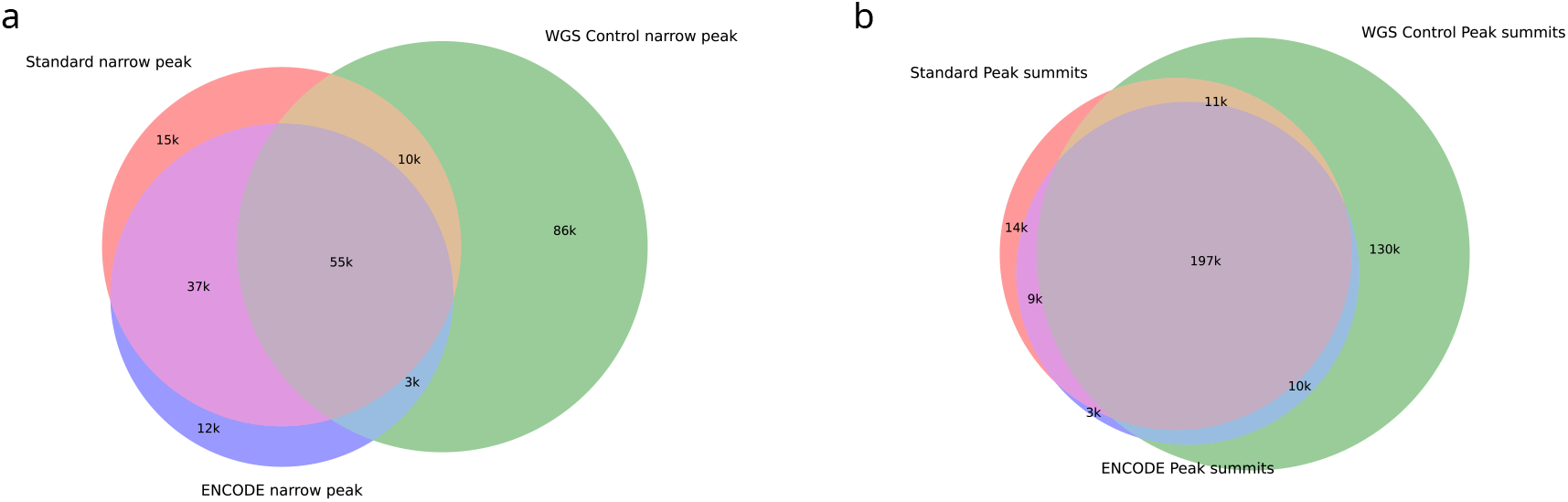
**a** Venn diagram of WGS-controlled peaks, standard peaks, and ENCODE peaks (reference). **b** Venn diagram of WGS-controlled peak summits (±500 bp), standard peak summits (±500 bp), and ENCODE peak summits (±500 bp).

**Fig. S3:**
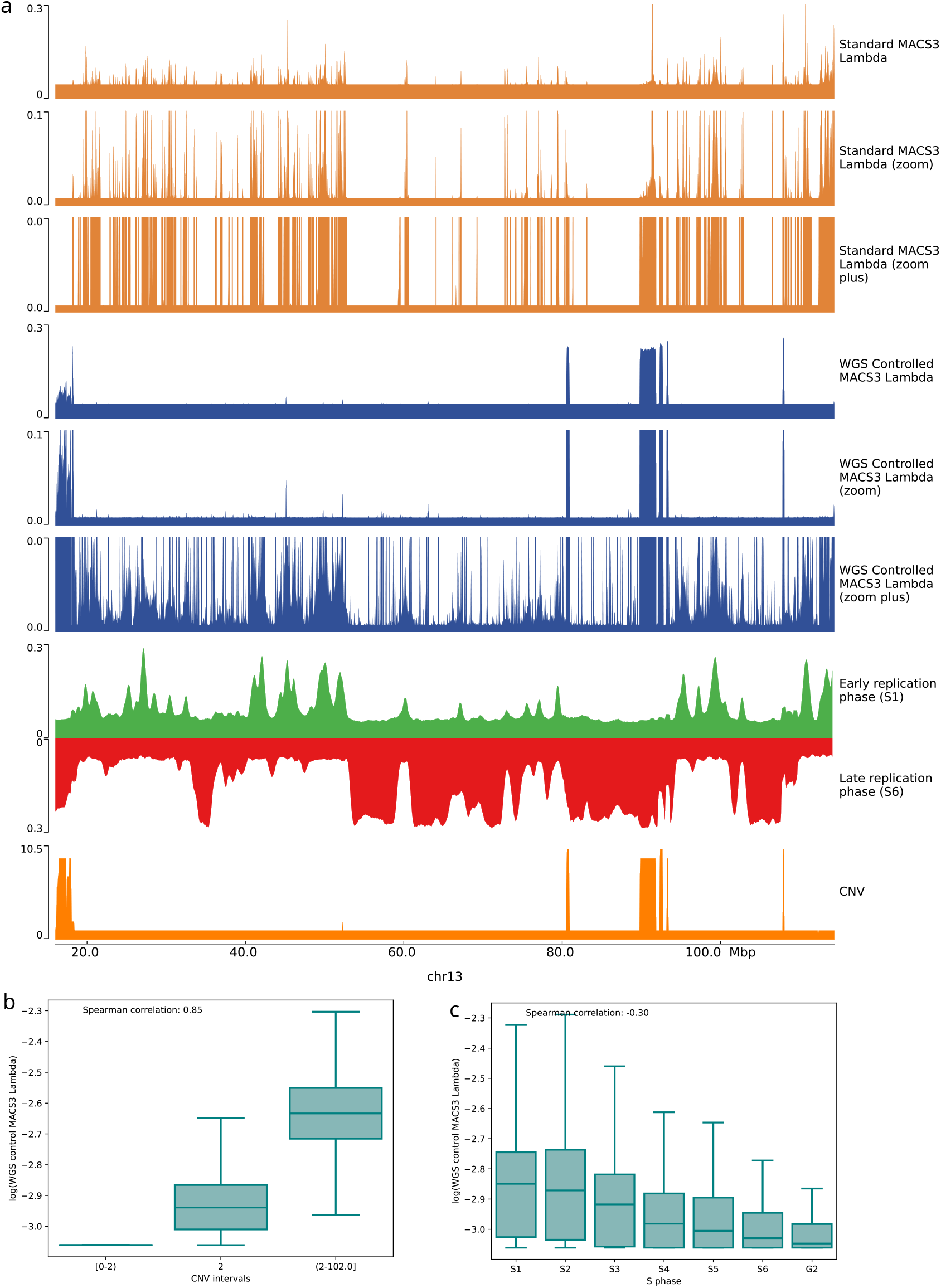
**a** Track view of standard ATAC-seq MACS3 lambda signal (orange), WGS-control ATAC-seq MACS3 lambda signal (blue), replication timing signal (S1 green, S6 red), and CNV (orange) on chromosome 13. ATAC-seq lambda signals are shown at three different scales. **b** Correlation between the MACS3 lambda computed with the peak-calling pipeline integrating WGS and CNV. The Spearman correlation between MACS3 lambda signal and mean CNV (1000-bp bins) is *ρ* = 0.85. **c** Correlation between the MACS3 lambda computed with the peak-calling pipeline integrating WGS and replication timing. The Spearman correlation between MACS3 lambda signal and replication timing phase is *ρ* = − 0.30, with a signal that decreases across replication timing phases.

**Fig. S4:**
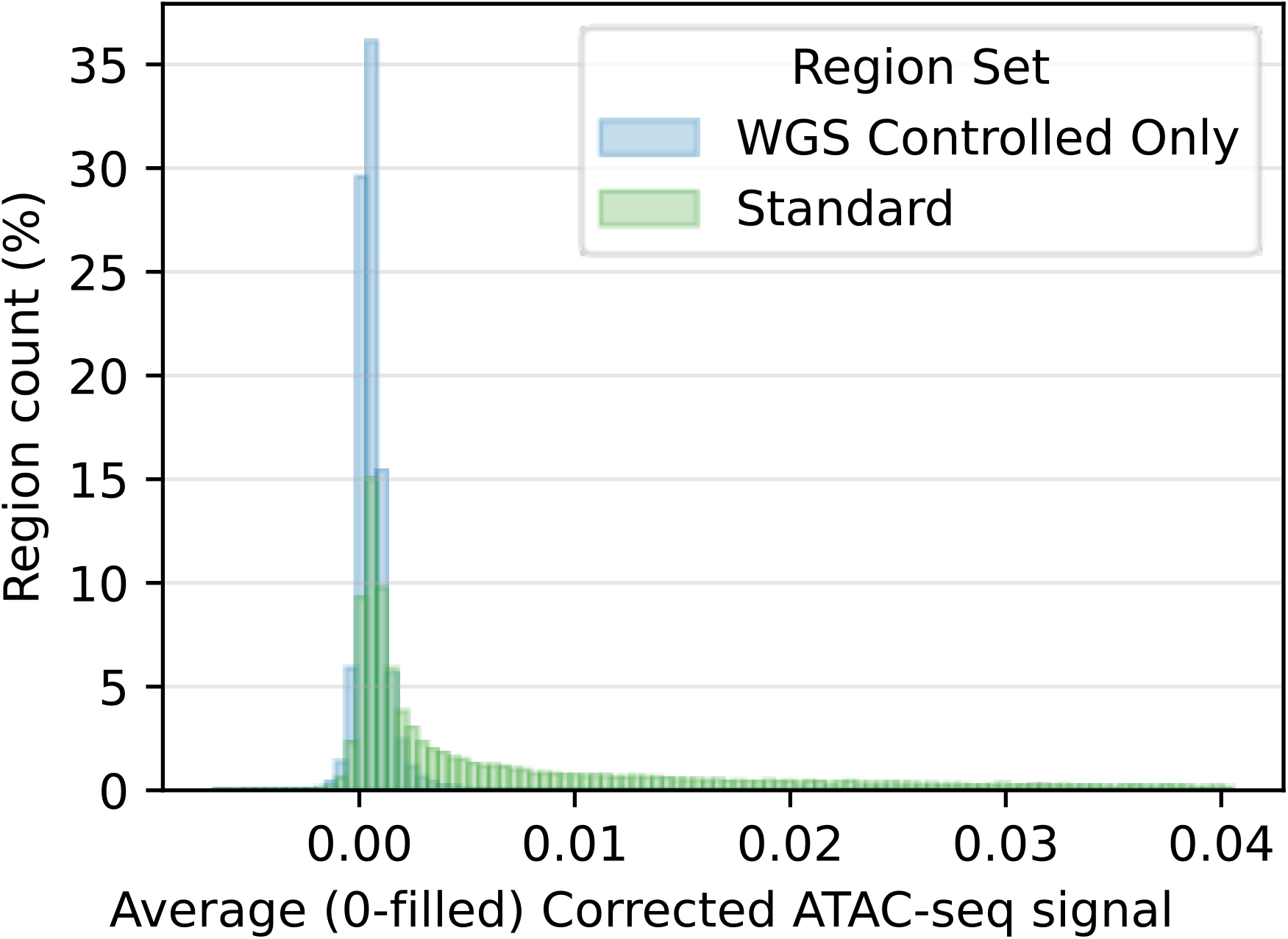
ATAC signal corrected for the Tn5 bias computed with TOBIAS `ATACorrect` on merged ATAC-seq BAM in WGS-controlled and standard peaks.

**Fig. S5:**
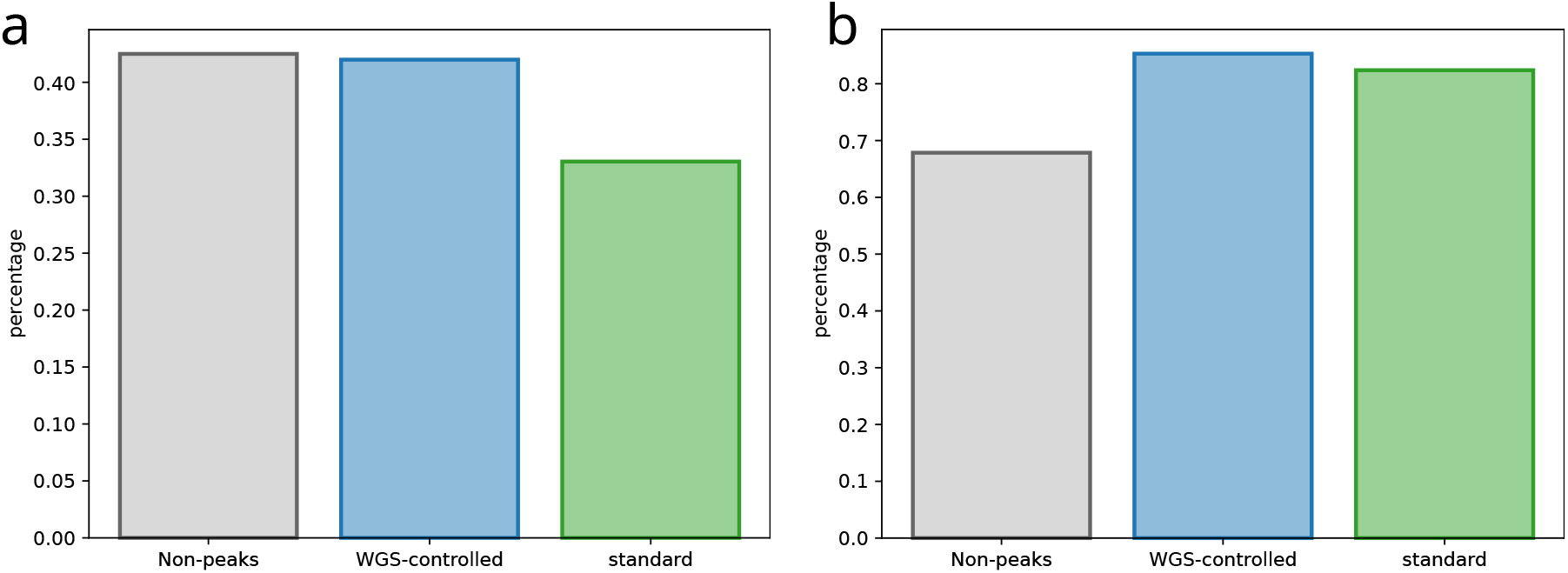
**a** Percentage of WGS-controlled (blue) and standard (green) peaks (summits ± 500bp) overlapping Alu repeat elements (respectively 41.9% vs 33.0%; Fisher exact test *p* = 0.0). Percentage of non-peak regions (grey, see Materials and Methods section) intersecting with Alu repeats is also shown. **b** Percentage of WGS-controlled (blue) and standard (green) peaks (summits ± 500bp) containing at least one eQTL (respectively 85.3% vs 82.4%; Fisher exact test *p* = 6.6 × 10^−133^). Percentage of non-peak regions (grey, see Materials and Methods section) containing eQTLs is also shown for comparison.

**Fig. S6:**
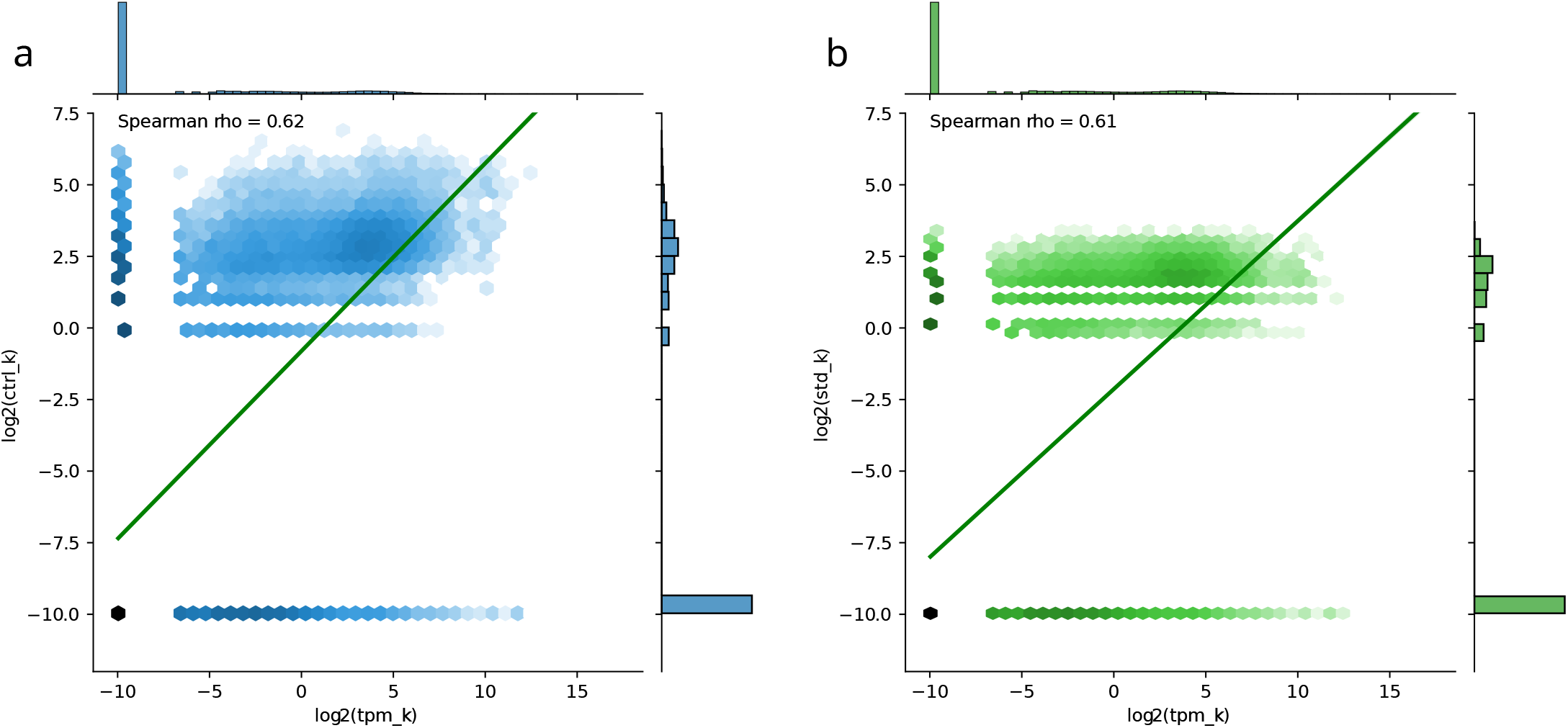
**a–b** Hexbin plots showing the correlation of the number of ATAC-seq peaks within a 2kb window centered around gene starts (y-axis) and gene expression (x-axis). ATAC peaks were called with WGS-integrating (a) and standard (b) pipeline. Spearman correlation *ρ* = 0.62 and 0.61 respectively.

**Fig. S7:**
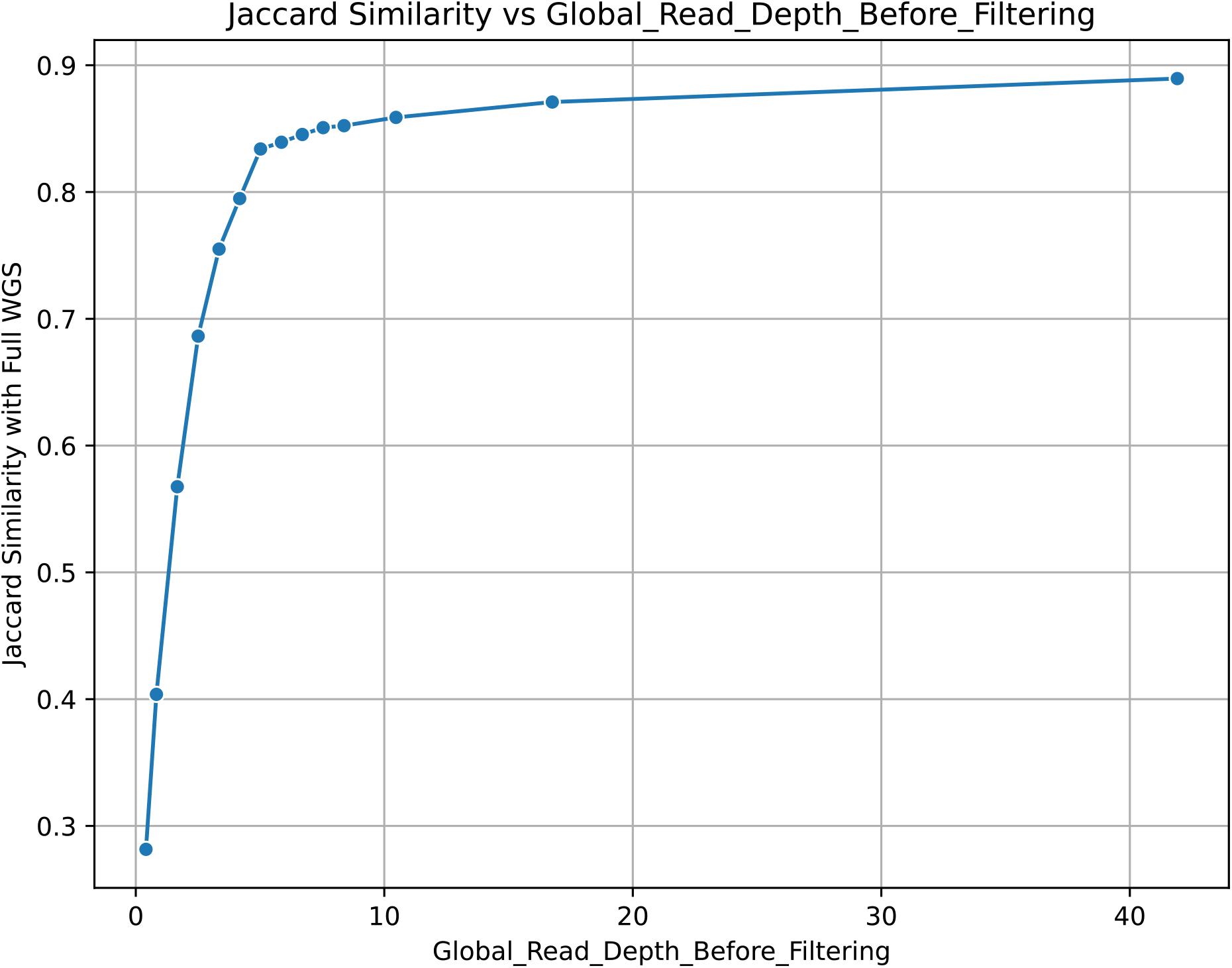
WGS FASTQ files were down-sampled by a factor of 0.5, 0.2, 0.125, 0.1, 0.09, 0.08, 0.07, 0.06, 0.05, 0.04, 0.03, 0.02, 0.01, and 0.005. At each iteration, the Jaccard index between the peaks called with and without down-sampled WGS data was computed. The Jaccard index drops when read depth is lower than 5 (down-sampling rate *<* 0.06 relative to original FASTQ files).

**Supplementary Table S1:**
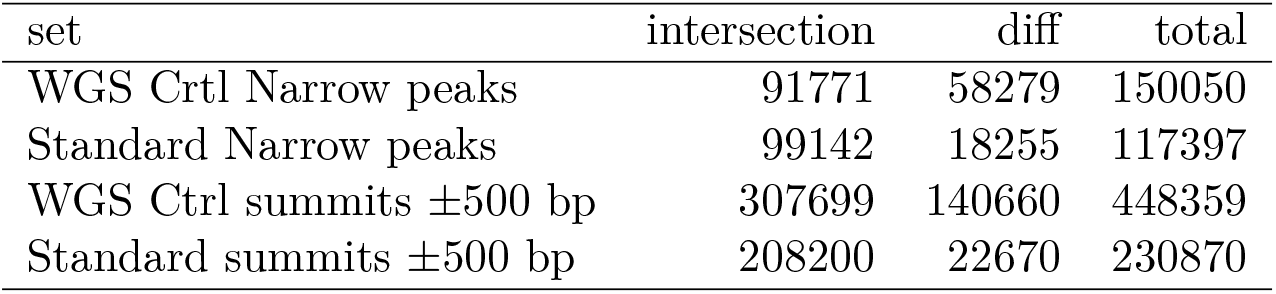
Intersection of peaks and peak summits called with WGS-integrating and standard pipelines.

| set | intersection | diff | total |
| --- | --- | --- | --- |
| WGS Ctrl Narrow peaks | 91771 | 58279 | 150050 |
| Standard Narrow peaks | 99142 | 18255 | 117397 |
| WGS Ctrl summits $\pm 500$ bp | 307699 | 140660 | 448359 |
| Standard summits $\pm 500$ bp | 208200 | 22670 | 230870 |

**Supplementary Table S2:** Peak length summary statistics.

| set | mean | median | 25th percentile | 75th percentile |
| --- | --- | --- | --- | --- |
| WGS Ctrl Narrow peaks | 624.448307 | 496 | 326 | 761 |
| Standard Narrow peaks | 375.777023 | 334 | 234 | 462 |

**Supplementary Table S3:** Fractions of peaks (summit ± 500bp) overlapping Alu repeats and eQTLs.

| Type | Intersect_WGS | Total_WGS | Fraction_WGS | Intersect_standard | Total_standard | Fraction_standard |
| --- | --- | --- | --- | --- | --- | --- |
| Alu | 59060 | 140660 | 0.4198777193231907 | 101674 | 307699 | 0.33043 |
| eQTLs | 119982 | 140660 | 0.8529930328451585 | 253486 | 307699 | 0.82381 |

**Supplementary Table S4:** ATAC-seq FE signal in WGS-controlled and standard peaks (summit ± 500bp).

| set | mean | median | 25th percentile | 75th percentile |
| --- | --- | --- | --- | --- |
| WGS-controlled | 0.1180 | 0.0989 | 0.0789 | 0.1319 |
| standard | 0.3816 | 0.1768 | 0.1079 | 0.3806 |

**Supplementary Table S5:** STARR-seq signal in WGS-controlled and standard peaks (summit ± 500bp).

| set | mean | median | 25th percentile | 75th percentile |
| --- | --- | --- | --- | --- |
| WGS-controlled | 1.5166 | 1.5166 | 1.0301 | 1.8767 |
| standard | 1.9268 | 1.7195 | 1.2220 | 2.3870 |

**Supplementary Table S6:**
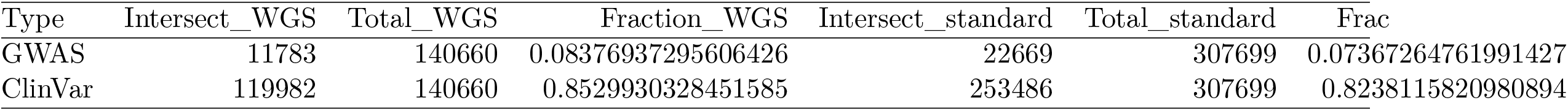
Fractions of peaks (summit ± 500 bp) overlapping GWAS causal traits and ClinVar variants.

**Supplementary Table S7:**
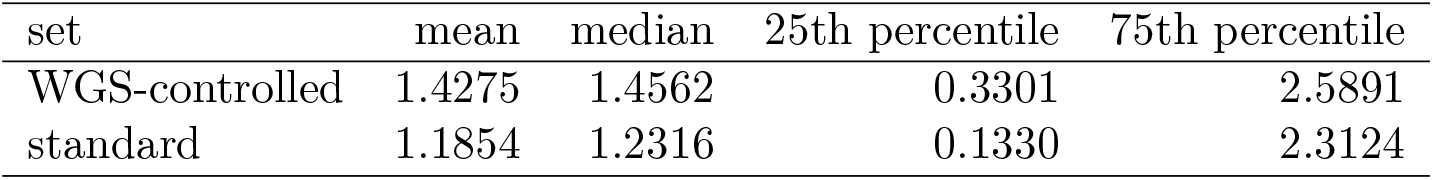
GnomAD score in WGS-controlled and standard peaks (summit ± 500bp).

| set | mean | median | 25th percentile | 75th percentile |
| --- | --- | --- | --- | --- |
| WGS-controlled | 1.4275 | 1.4562 | 0.3301 | 2.5891 |
| standard | 1.1854 | 1.2316 | 0.1330 | 2.3124 |

**Supplementary Table S8:**
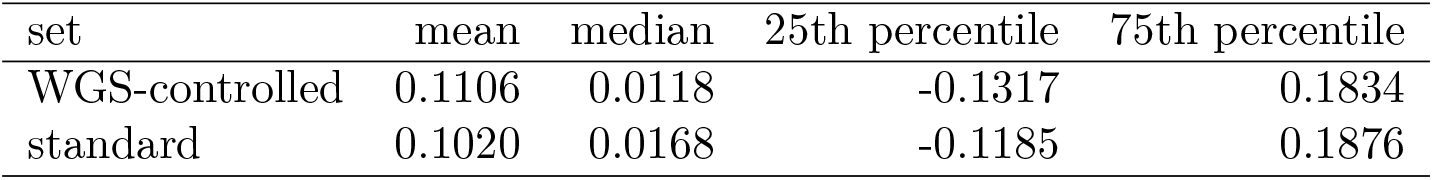
phyloP score in WGS-controlled and standard peaks (summit ± 500bp). set mean median 25th percentile 75th percentile.

| set | mean | median | 25th percentile | 75th percentile |
| --- | --- | --- | --- | --- |
| WGS-controlled | 0.1106 | 0.0118 | -0.1317 | 0.1834 |
| standard | 0.1020 | 0.0168 | -0.1185 | 0.1876 |

**Supplementary Table S9:** Fraction of WGS-controlled peaks (all peaks summit ± 500bp) called in K562 intersecting with MCF-7 WGS-controlled peaks (all peaks summit ± 500bp) and fraction of standard peaks (all peaks summit ± 500bp) called in K562 intersecting with MCF-7 standard peaks (all peaks summit ± 500bp).

| Intersect_WGS | Total_WGS | Fraction_WGS | Intersect_standard | Total_standard | Fraction_standard |
| --- | --- | --- | --- | --- | --- |
| 58571 | 140660 | 0.416401 | 195172 | 307699 | 0.634295 |

